# Novel Single-Cell Multiomics Approach to Analyze Replication Timing and Gene Expression in Mouse Preimplantation Embryos

**DOI:** 10.1101/2025.01.02.631157

**Authors:** Anala V. Shetty, Clifford J. Steer, Walter C. Low

## Abstract

Analysis of a cell’s replication timing (RT) provides insight into how genes replicate, early or late, during the S-phase of the cell cycle. RT is cell-type specific, inheritable, and has been correlated to gene expression in normal and diseased states. However, most studies have been limited to somatic cells. Very little is known about RT control in early mouse embryos, and how it correlates with the start of transcription during zygote gene activation (ZGA), at the 2-cell stage. In this study, we developed a novel in-house single-cell multiomics approach to simultaneously analyze RT and gene expression in individual cells of the mouse 1-cell, 2-cell, and 4-cell embryos. We detected that RT was established at the 1-cell stage prior to ZGA. Surprisingly, we observed that the coordinated RT and gene expression control was different in early totipotent embryos, compared to previously published studies in somatic cells. Late replicating regions correlated with higher gene expression and open chromatin in the early developing embryos. Lastly, we performed an integrated pseudo time trajectory analysis combining RT and gene expression information per cell.

## Introduction

Over the past 20 years, replication timing (RT) of cells has been studied to understand how cells replicate their genome during the S-phase of the cell cycle and how these coordinates with genome organization and function^1–4^. RT changes occur in ranges of 400-800 kbps, are independent of the DNA nucleotide sequences, and conserved within different cell types^5^. RT has been correlated with gene expression, chromatin accessibility, and positioning inside the nucleus, making RT an epigenetic regulator^2,6–11^. Multiple studies have demonstrated how RT is disrupted in disease states such as physiological and premature aging^1,12^, down syndrome^13^, and various types of cancer^14,15^. Most of these RT studies have been limited to somatic cells. Very few studies have examined at RT in totipotent cells of the early mouse embryo^16,17^. Recently, we developed an affordable in-house single cell (sc) multiomics approach that allows us to simultaneously extract and process both gDNA and mRNA from each single cell. This protocol is compatible with very small starting cell numbers (as low as 10 cells), making it ideal for processing the early embryos. It is also compatible with both cells and nuclei. Briefly, gDNA and mRNA were extracted and separated from each cell using the Smart-seq2 technique,^18^ where magnetic beads with poly dT tailed primers were used to pull down the poly A tailed mRNA. The gDNA was amplified using an in-house protocol, the mRNA was also processed in house as described in the G&T protocol.^19^ Sc-RT and gene expression information were derived from the amplified gDNA and mRNA respectively. Further, we could also make high precision correlations between the two parameters within each cell.

In this study, we used the sc-multiomics protocol to analyze cells from the earliest moue embryonic stages-zygotes, 2-cell, and 4-cell stages (**Fig. 1**). We were able to (i) analyze sc-RT and sc-RNA patterns at the 3 stages; (ii) draw novel correlations between RT and gene expression within each stage; (iii) analyze genome-wide trends in RT, gene expression, and chromosome accessibility changes between developmental stages; and (iv) perform a novel integrated pseudo time trajectory analysis using both scRT and scRNA data.

**Figure 1:**
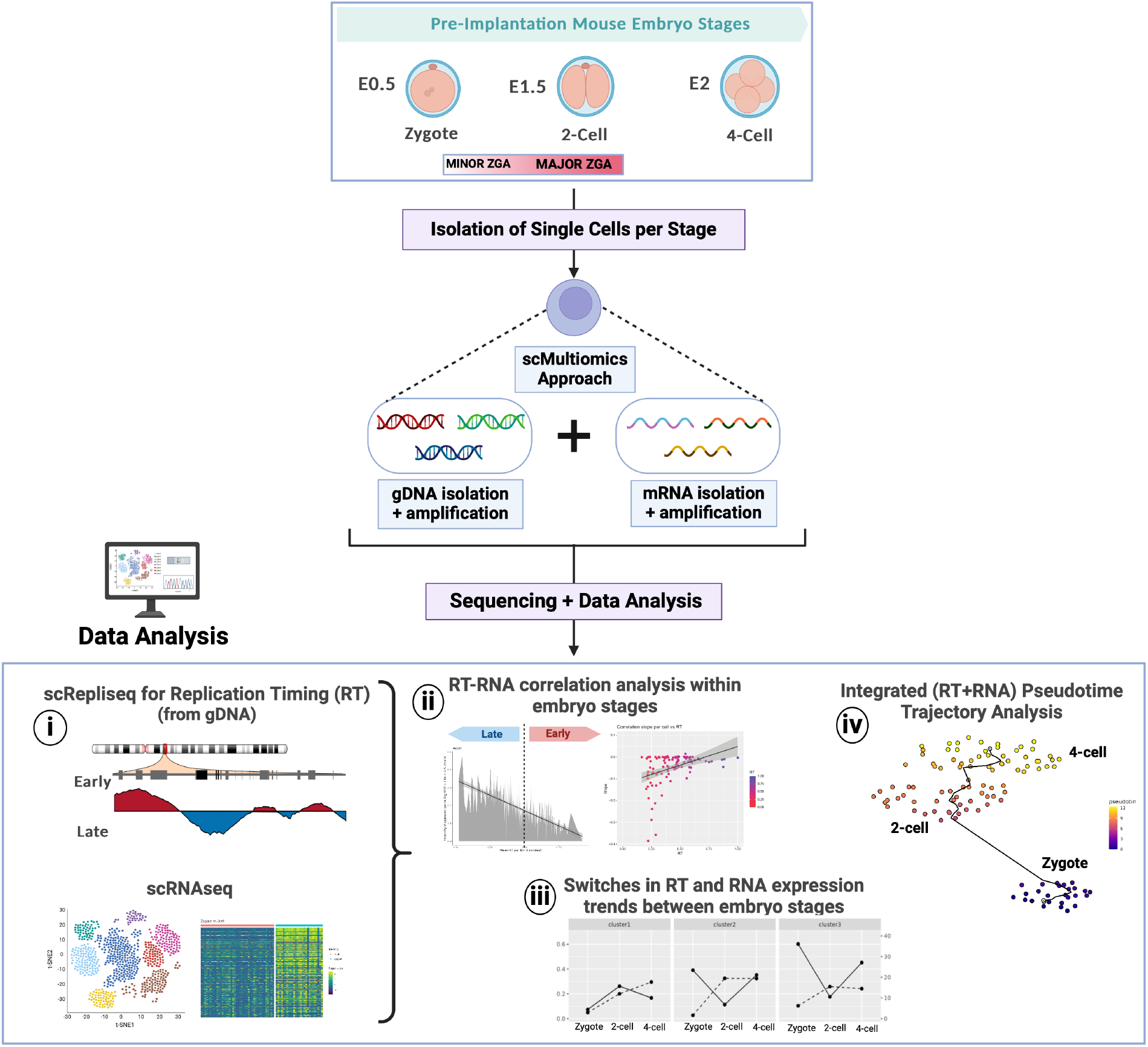
Novel Single Cell-Multiomics Approach to Study Replication Timing and Gene Expression in Pre-Implantation Embryos. Single cells were extracted from zygotes, 2-cell and 4-cell embryos and processed our novel in-house single cell multiomics approach for parallel analysis of RT and gene expression. The main results of the analysis done are depicted (1-5). **(i)** gDNA was used to deduce single cell Replication Timing (sc-RT) trends. Gene expression information was obtained from the sequenced mRNA. **(ii)** Correlations between RT and gene expression was performed within each embryo stage and within individual cells. **(iii)** Switches in RT and gene expression trends between the 3 developmental stages was analyzed. **(iv)** A novel integrated pseudo time trajectory analysis using both RT and gene expression per cell was performed.

Very little is known about RT at the earliest embryonic stages and if RT is established immediately after fertilization. Recently, two groups analyzed sc-RT in early mouse embryos^16,17^, however, correlations to gene expression within the same cells, and at the single cell level, have not been done to date. An important question in the RT field is whether RT precedes gene expression or vice versa. Major transcription does not begin in embryos until the 2-cell stage^20^. Hence, analyzing the zygote stage will allow us to conclude if there is a stable RT at the zygote stage, or if transcription is required for establishment of RT.

The embryos at these early stages are totipotent and have unique nuclear organization compared to somatic cells. The early embryos undergo rapid genome restructuring in the nucleus ^21^. At the zygote stage, the maternal and paternal chromosomes are separate clusters present in pronuclei, within the same cytoplasm^22^. The paternal chromatin, which is highly compressed in the sperm, undergoes rapid decompression in the zygote^22,23^. Until the 4-cell stage, both sets of chromosomes are not completely mixed within the nucleus^21^. The somatic cell types studied to date have a stable nuclear organization where the DNA is tightly packed into chromatin onto histones^24^. The topological boundaries remain intact, and any rearrangements, such as during stem cell differentiation, occur at a much smaller scale compared to early embryos^25^. These features, that are unique to the rapidly replicating mouse embryo, make it a unique model to study developmental RT and correlate it to gene expression using the sc-multiomics approach.

### Section 1 Replication timing was established at the zygote (1-cell) stage and was further defined as embryonic development progresses to 2-cell and 4-cell stages

Single cell RT was analyzed for early preimplantation mouse embryos at zygote/1-cell, 2-cell, and 4-cell stages. Single cells from these stages were processed using our in house sc-multiomics approach and analyzed using the Kronos scRT pipeline^26^ as described in the methods. Single cell gDNA was used to derive sc-RT profiles.

**Fig. 2a** illustrates the distribution of cells across the early, mid, and late S-phase for zygotes (n= 36), 2-cell (n=42) and 4-cell (n=43) stages. Each cell is plotted with its percentage of replicated genome plotted on the Y axis. The sc-RT profiles of S-phase cells, for chromosomes 2 and 3, are shown in **Fig. 2d**. The pseudo bulk RT profiles for each chromosome are shown in green above each sc-RT plot.

**Figure 2:**
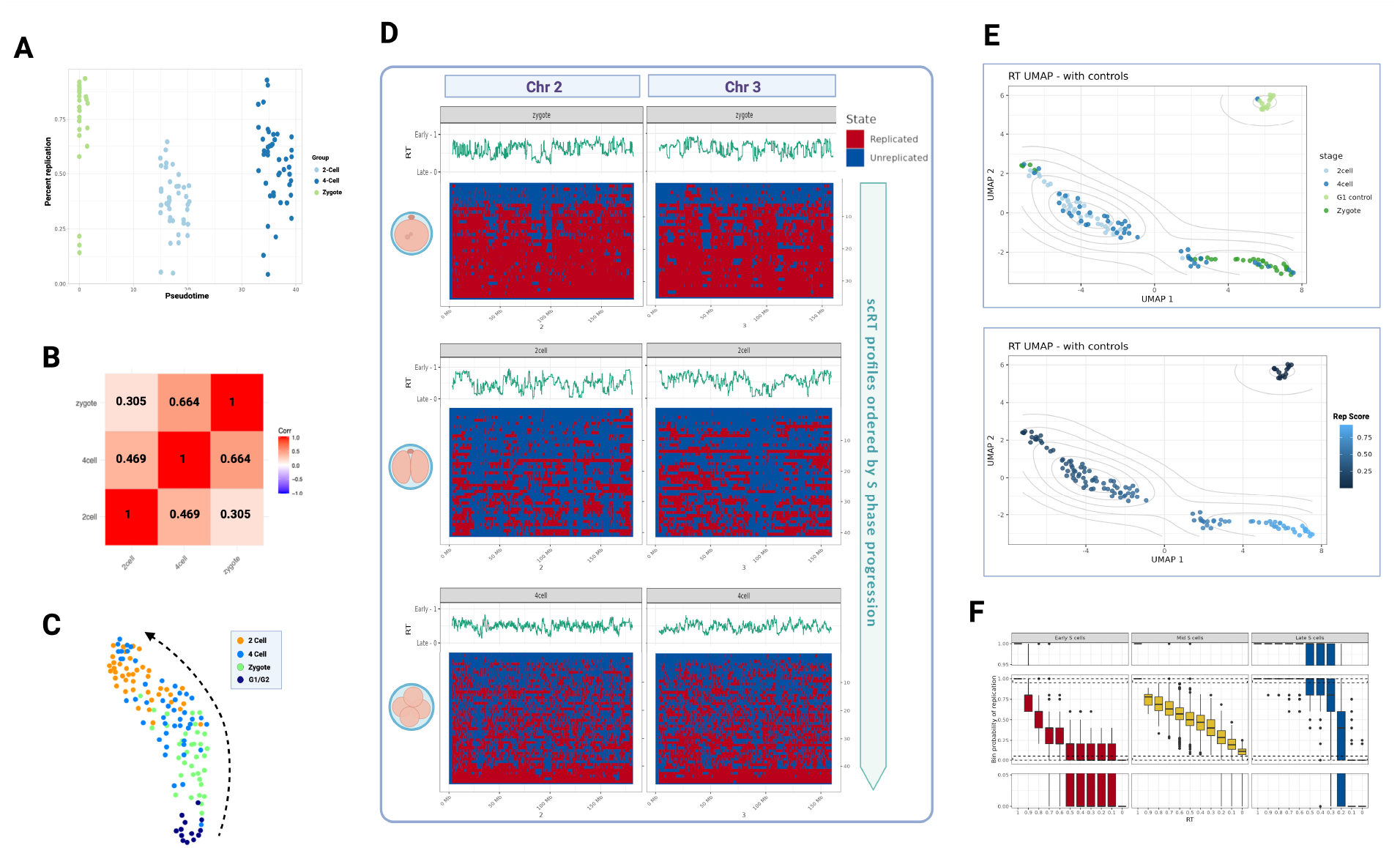
Replication Timing is Established at the 1-Cell Stage and is Further Defined as the Embryo Develops. **(a)** Percentage of genome replicated in individual S-phase cells analyzed from zygotes (n=36), 2-cell (42) and 4-cell embryos (n=43). gDNA from these cells was used for generating scRT trends. **(b)** Correlation of pseudo bulk RT between zygotes, 2-cell, and 4-cell embryos. **(c)** SPRING algorithm used for visualizing RT trajectory across single cells from G1/G2 mouse embryonic stem cells (used as controls), zygotes, 2-cell, and 4-cell embryos. **(d)** Chromosome 2 (chr2) and chromosome 3 (chr3) single cell RT binarized plots for zygotes (top panel), 2-cell (middle panel), and 4-cell (bottom panel) embryos. Cells are ranked by S-phase progression. Each row is a single cell RT profile where the red bins are replicated regions, and the blue bins are non-replicated regions. The pseudo bulk trends are in green above each binarized plot. **(e)** Single cell RT based UMAPs generated using G1/G2 control cells, zygote, 2-cell, and 4-cell stages. Top panel shows the cell populations and bottom panel shows the Rep Score, that is, progress through S-phase, for each cell. **(f)** The bin probability of replication in early S, mid S and late S phase, for bins genome-wide, are shown for the 4-cell stage. This was an output as part of the Kronos scRT pipeline that was used for generating scRT profiles.

In the zygote sc-RT profiles (**Fig. 2d (top panel**)), we observed common patterns of late replicating blue bins that were conserved across the S phase single cells. From those analyzed, we observed early replicating zygotes with <20% replicated genome, and late replicating zygotes with >57% of the genome replicated (**Fig. 2a)**. Very few mid-S phase zygotes with ∼ 40-50% of the genome replicated were present. The absence of mid-S phase zygotes was not due to processing time as all zygotes were lysed within 1 hour of extraction. Rapid switching from non-replicated to ∼60% replicated might be a feature of this earliest embryonic stage where replication takes place in simultaneous large bins across the genome. To confirm that the replication pattern that we observed for the zygotes was indeed defined and not random, we compared their pseudo bulk RT profile with RT profiles from two reference embryo datasets recently published^16,17^. Approximately 70% of total bins across the genome were conserved between all three RT profiles (**Extended data Fig. 1b**). This underscores that conserved RT patterns are established at the 1-cell stage.

We also observed RT patterns across single cells in 2-cell and 4-cell stage embryos. As the embryos progressed through the 2-cell and 4-cell stages, the size of early and late replicating domains got smaller and more defined (**Fig. 2d)**. Single cells analyzed from the 2-cell embryos were distributed in the mid-S phase (**Fig. 2a**) where approximately half the genome was replicated. Replicated and non-replicated regions of the genome corresponded to early and late replicating bins. Conserved RT patterns could be observed across the sc-RT profiles in **Fig. 2d**. For the 4-cell stage, a demonstrable distribution of cells in the early, mid, and late S-phase was observed (**Fig. 2a**). The replication bins were even smaller and more defined than the 2-cell embryos (**Fig. 2d)**. Gradual progression from early to mid to late S phase cells could be observed. The probability of replication in different bins across the genome, for early, mid, and late S phase cells could be observed for the 4-cell stage embryo in **Fig. 1f**. The number of bins replicating in the early, mid, and late S-phases could be observed from the per stage Twidth plots and the S phase window in which cells from each stage replicate can also be observed in **Extended data Fig. 1a**.

Moderate Pearson’s correlation was observed between RT profiles for the zygote, 2-cell and 4-cell stages (**Fig. 2b**). In the RT UMAP in **Fig. 2e**, zygotes, 2-cell, and G1 mouse embryonic stem cells (G1 controls) formed separate clusters. Single cells from the 4-cell embryos were distributed between the zygote and 2-cell clusters. Within the G1 cluster, we could also observe G1 single cells from the 2-cell and 4-cell stages. Using the SPRING algorithm^27^, we were able to visualize the RT trajectory across the developmental stages (**Fig. 2c)**. The 4-cell stage cells, similar to the RT UMAP, were distributed between zygotes and 2 cells.

### Section 2 Transcriptome profiles recapitulated stage-specific gene expression

From the same set of embryo cells used for the RT analysis, mRNA was extracted, processed, and analyzed as described in the methods. Unlike somatic cells where transcription is always present during different phases of the cell cycle, it is absent in the early embryo until the 2-cell stage. At the early 2-cell stage, a burst of transcription occurs through the major zygote gene activation (ZGA) process^28^. Recent studies have demonstrated that a minor ZGA, with very low levels of transcription, begins at the late zygote stage^20^. The 3 embryonic stages clustered separately on the gene expression UMAP (**Fig. 3a**), demonstrating stage-specific gene expression. We were able to visualize the progression through the embryonic stages using pseudo time trajectory analysis (**Fig. 3c**). We observed a high correlation in gene expression between our sc-RNA datasets ^29,30^and previously published reference datasets (**Extended data Fig. 1c**).

**Figure 3:**
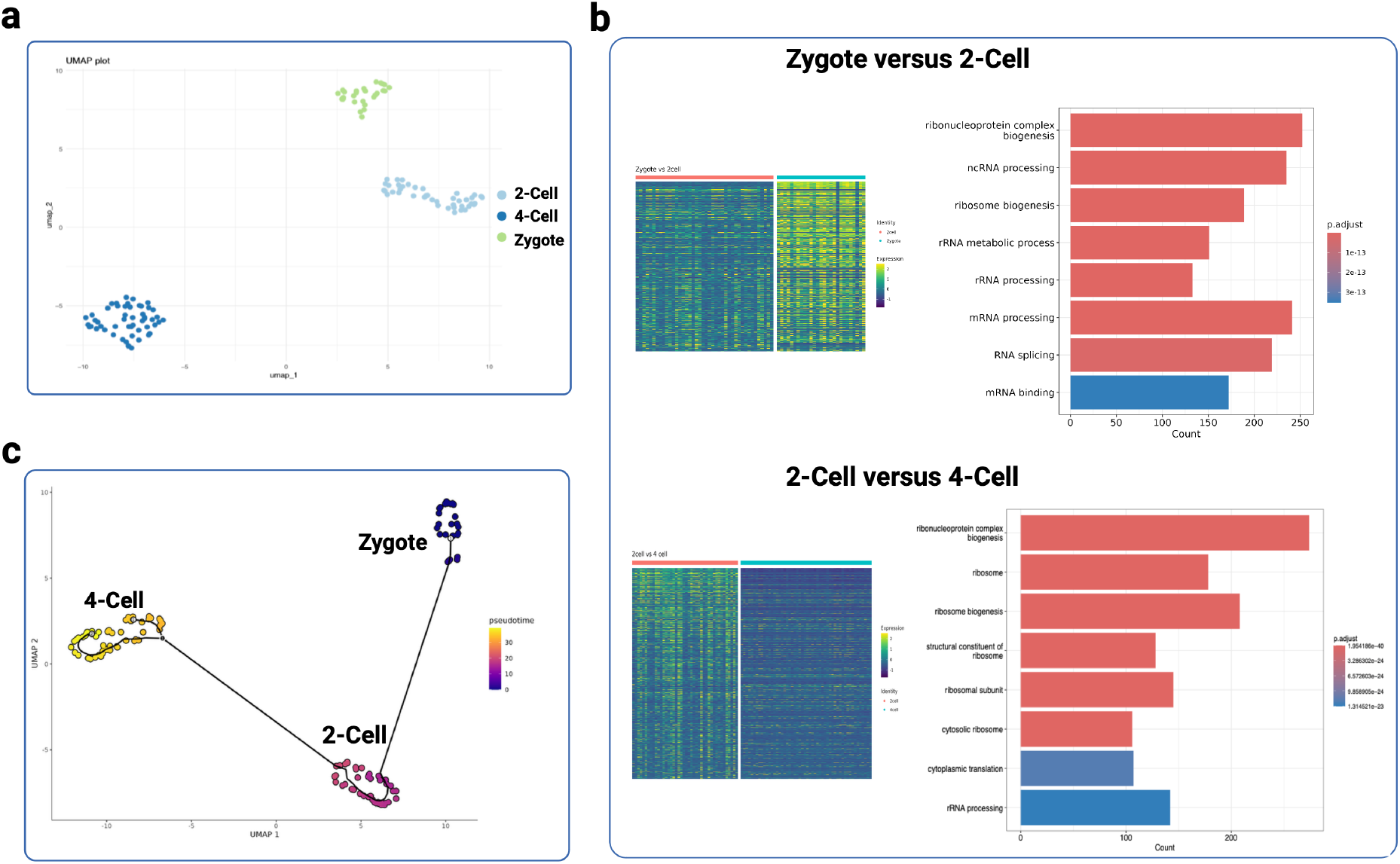
Gene Expression Analysis of Single Cells from Zygote, 2-cell, and 4-cell Embryos. **(a)** Gene expression UMAP depicting stage-specific clusters for zygote, 2-cell, 4-cell embryos. **(b)** Pairwise comparison of gene expression between zygote and 2-cell (top panel) and 2-cell and 4-cell (bottom panel) embryos. Heatmaps of top differentially expressed genes (DEGs) and the corresponding Gene Ontology terms of the top DEGs. **(c)** Pseudo time trajectory analysis of zygote, 2-cell and 4-cell embryo cells.

In addition, gene ontology of top differentially expressed genes between zygote and 2-cell, and 2-cell and 4-cell stages highlighted activities related to RNA processing and splicing (**Fig. 3b** top panel) and ribosome processing and translation (**Fig. 3b** bottom panel).

### Section 3 Higher levels of gene expression occur in late replicating bins in 2-cell and 4-cell embryos

Multiple replication timing-gene expression (RT-RNA) correlation studies have been reported. However, the majority of these studies were performed in somatic cells, including stem cells, and differentiated cell types^2,31–33^. Further, the studies to date used separate populations of cells for processing RT and gene expression. In this study, extracting both gDNA and mRNA from the same cells enabled us to perform high precision, genome-wide correlations between RT and gene expression within individual cells, which to our knowledge, has not been previously reported.

First, we performed a pseudo bulk analysis of the RT-RNA correlation for cells from zygote, 2-cell, and 4-cell stages to understand the general trend observed within each stage. We used the stage-specific pseudo bulk RT profile and considered gene expression across all cells as described in the methods. We plotted the probability of gene expression against genome wide RT bins, considering the total transcriptome per stage. However, with the total transcripts, we observed no preference of early or late bins for gene expression (**Extended Data Fig. 2a**). The same lack of preference was also previously observed when total transcriptome was correlated with RT in the zygotes, 2-cell, and 4-cell, from separate datasets of sc-RNA and sc-RT^16^. However, at the earliest embryonic stages, maternally inherited transcripts from the oocyte are present in the embryos^34^. These maternal mRNAs (i) make up the majority of total transcripts at the earliest stages; (ii) are not actively transcribed in the embryo; and (iii) can skew the correlation between gene-specific RT and stage-specific transcription of those genes. Hence, we re-analyzed the RT-RNA correlation, considering only post-ZGA transcripts for each stage, subtracting out the maternal mRNAs as described. For the post-ZGA gene list, we observed a RT-RNA correlation. Higher gene expression was more prominent in the late replicating bins compared to the early replicating bins in 2-cell and 4-cell stages (**Fig. 4a, Fig. 4b**). For the zygote stage, this trend was less significant as very few genes are transcribed at this stage.

**Figure 4:**
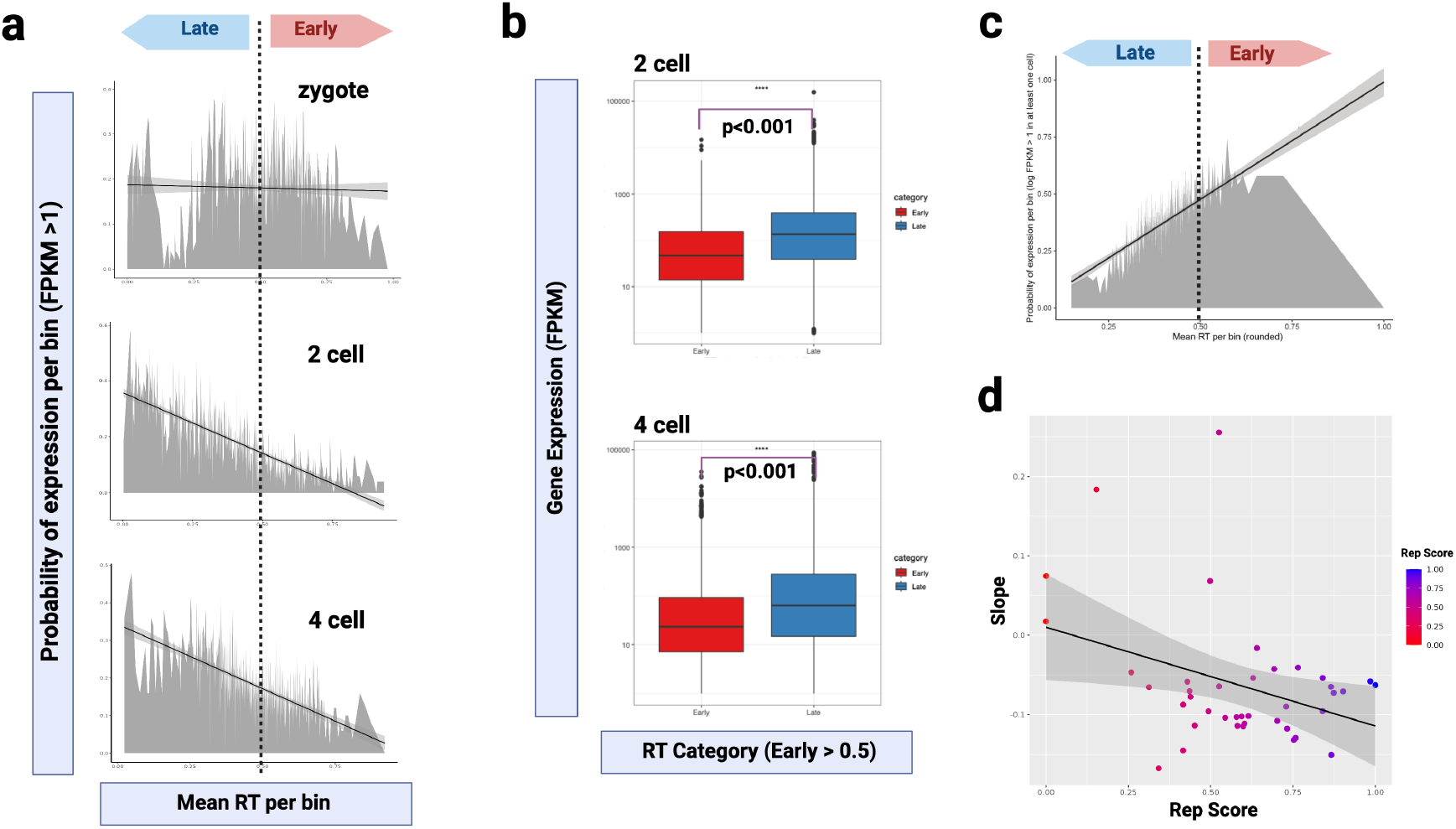
Correlation Trends between Replication Timing and Gene Expression in Embryos Differs from Somatic Cells. **(a)** Pseudo bulk analysis of correlation between RT and gene expression within each stage. The probability of gene expression per bin was plotted against gene-specific RT bins (50 genes per bin) genome-wide. Late (RT <0.5) and early (RT>= 0.5) are depicted in Zygote (top panel), 2-cell (middle panel), and 4-cell (bottom panel) embryos. The RT-gene expression correlation slope (negative slope) is also depicted for each stage. **(b)** The gene expression in late versus early bins for 2-cell and 4-cell stages are depicted. **(c)** The pseudo bulk analysis for HepG2 cells processed using our in-house sc-multiomics protocol. The RT-gene expression correlation slope (positive slope) is depicted. **(d)** RT-gene expression correlation trends for individual cells analyzed from the 2-cell embryo stage. The RT-gene expression correlation slope was calculated per cell and plotted on the Y axis. The Rep Score, that is, percent of genome replicated, is plotted on the X axis.

This preference of late replicating bins for higher gene expression contrasts with what has been observed for somatic cell types^2,35,36^. In more differentiated cell types, higher gene expression was more prominent in the early bins compared to the late bins. We performed the RT-RNA correlation for the human HepG2 cell line, which is a somatic cell line, processed in-house using the same sc-multi omics protocol. We observed a trend similar to previously published studies on somatic cells, observing higher gene expression in early replicating bins compared to late bins (**Fig. 2C**). The contrasting trend observed in embryos could be due to the rapid reorganization taking place until the 4-cell stage^24^. In somatic cells, since chromatin exists in a more stable repressive conformation, replication mechanisms are characteristically different than that of the embryo.

We also performed RT-RNA correlations within each individual cell as described in the methods. Similar to the negative slope trend of the 2-cell pseudo bulk analysis, 37 out of 42 cells from the 2-cell stage had a negative slope value implying higher gene expression in late replicating bins within each cell (**Fig. 2d**). For the zygote stage, a low positive slope was observed in the individual cells (**Extended Data Fig. 2b**), like the zygote pseudo bulk trend. However, significant RT-RNA correlations could not be made due to the low number of transcripts at this stage. For the 4-cell stage, the very early (<25% genome replicated) and very late (>75% genome replicated) S phase cells had a positive slope value but cells in the mid-S phase were distributed between negative and positive slope gradients. The 4-cell pseudo bulk plot, generated from the same population of cells, had a significantly negative slope (**Extended Data Fig. 2c**) which was not conserved in all the individual cells from the 4-cell stage. This highlights and underscores the importance of the sc-multiomics analysis in identifying individual cell trends which may not be reflected in pseudo bulk or bulk analyses.

### Section 4 Switches in RT Correlated with Chromosome Accessibility and Gene Expression

In the previous section, we looked at RT-RNA trends within each developmental stage. Next, we wanted to observe how RT trends changed between embryonic stages and its links to other genomic features. We performed genome-wide hierarchical clustering of RT bins, across the 3 stages (clusters 1-8), as described in the methods. Early-to-late or late-to-early RT switches between stages were observed (**Fig. 5a**) and GO of switching clusters are listed in **Extended Fig. 3c**. Within each cluster, we plotted the mean RT and gene expression and observed changes between embryonic stages.

**Figure 5:**
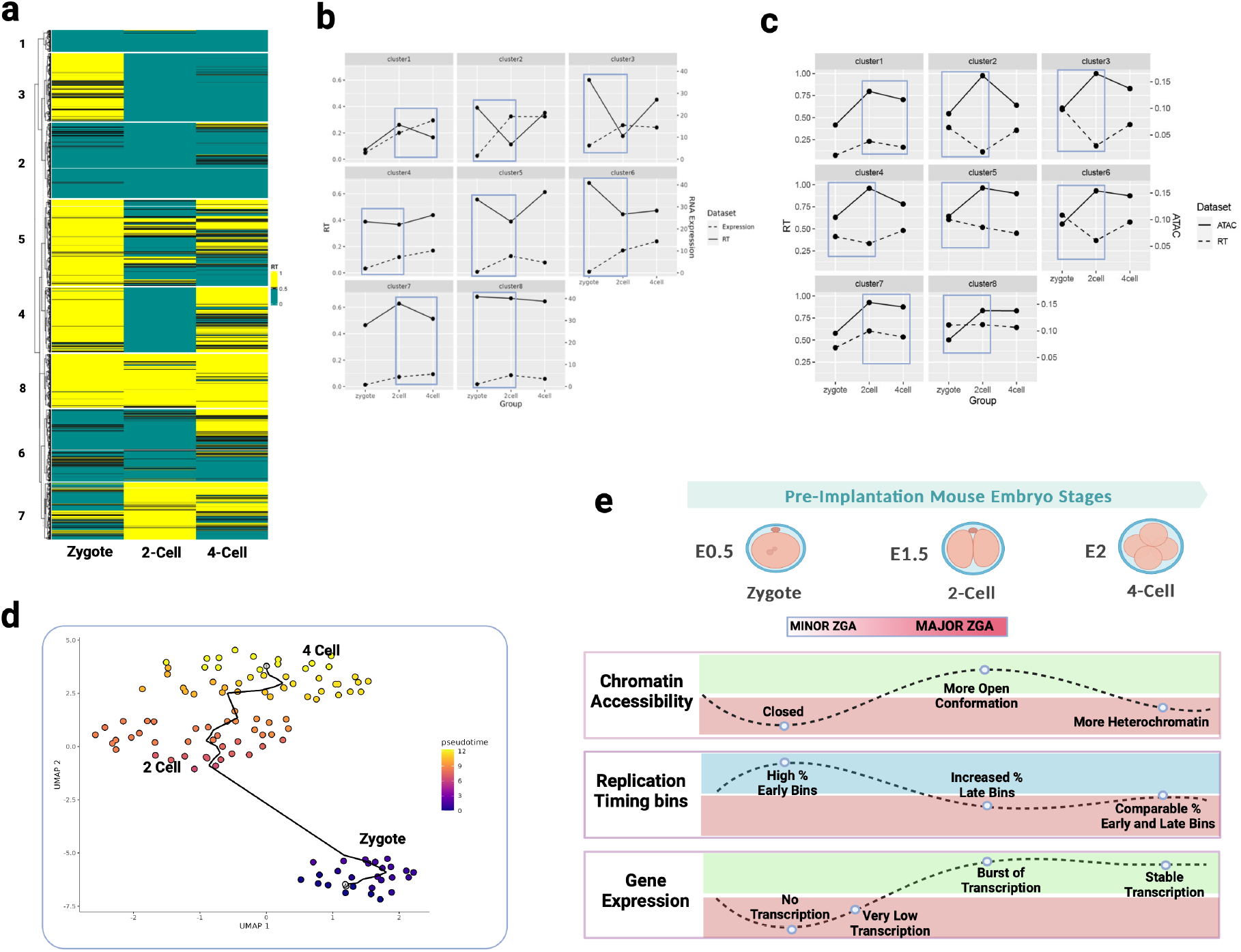
Developmental changes in Replication Timing, Gene Expression, and Chromatin Accessibility during Early Mouse Embryogenesis. **(a)** Hierarchical clustering of RT bins genome wide, across the 3 embryonic stages. The early replicating bins (RT values >=0.5) are shown in yellow, and the late bins (RT values <0.5) are shown in green. **(b)** Trends of changes in mean RT and gene expression for all clusters 1-8 from Figure 5A. Blue boxes highlight trends of late-to-early RT changes and corresponding lower-to-higher gene expression changes. **(c)** Trends of changes in RT and chromosome accessibility (ATAC-seq values) for all clusters 1-8 from Figure 5A. Blue boxes highlight trends of late-to-early RT changes and corresponding closed-to-open chromatin changes (higher ATAC score). **(d)** Integrated trajectory analysis of zygote, 2-cell, and 4-cell embryonic stages. Trajectory was based on integrated RT and gene expression information per cell. **(e)** Summary of chromosome accessibility, RT, and gene expression trends observed genome wide across zygotes, 2-cell, and 4-cell embryonic stages.

A common trend that was observed across all clusters was evidenced by earlier-to-later RT switches between stages which correlated with lower-to-higher gene expression (**Extended Data Fig. 3a**). This finding supported the higher gene expression in late bins compared to early bins result that we observed within stages (**Fig. 4a)**. We also observed that clusters that were constitutively early replicating across all stages (clusters 7 and 8), with mean RT >=0.5, had lower gene expression levels compared to clusters that were constitutively late replicating (clusters 1, 2). We did not observe genome-wide conserved trends of late-to-early RT switches and lower gene expression. The presence of transcripts in the cell, even after transcription is turned off, would make this difficult to quantify.

Next, we correlated chromatin accessibility at gene promoter regions with gene-specific RT and expression. We hypothesized that in the developing embryos, genes in late replicating bins have more open chromatin, making them accessible to the transcription machinery, and hence show higher gene expression. To test this, we integrated an ATAC-seq reference dataset for embryos (GSE66581) with the RT and RNA datasets generated in the study. We examined ATAC, RT, and RNA trends in clusters 1-8, considering post-ZGA genes present in all three datasets. An increase in ATAC value, which corresponded to more open chromatin conformation, correlated with an increase in gene expression in the embryos (**Extended Data Fig. 3b**), as previously demonstrated^29^. In ATAC trends across all clusters, an inverted ‘V’ trend was observed. The ATAC value increased from the zygote to the 2-cell before decreasing by the 4-cell stage. This indicated a closed conformation at the zygote stage, opening up at the 2-cell stage, and a more closed conformation at the 4-cell stage. These observations are in agreement with previous observations^29^.

Early-to-late RT switch correlated with lower-to-higher gene expression (**Fig. 5b**). Across all clusters, an increase in ATAC value correlated with early-to-late RT switches (**Fig. 5c**). In cluster 1, the mean ATAC value did not increase with the early-to-late RT switch but remained about the same. However, cluster 1 was very late replicating across all three stages and had progressively increased gene expression (**Fig. 5b**). Overall, considering genome-wide ATAC, RT, and RNA trends, a closed-to-open chromatin conformation correlated with early-to-late RT transition and lower-to-higher gene expression.

We observed a drastic increase in the number of genome-wide late bins from zygote to the 2-cell stage (**Extended Data Fig. 3d**). An increase in open chromatin, from zygote to the 2-cell stage, could also be observed (**Extended Data Fig. 3d**). These results suggested that the zygote stage chromatin becomes more accessible at the 2-cell stage. This drastic opening up of the 2-cell stage chromatin coincided with the major-ZGA when a burst of transcription is observed (**Fig 5e**). Previous studies have also demonstrated drastic rearrangement of chromatin at the 2-cell stage^22,37^.

In clusters 2-6, we also observed a late-to-early RT trend (**Fig. 5c**); and a corresponding open-to-closed ATAC trend (**Extended data Fig 3b**) was observed for 2-cell-to-4-cell transition. This suggested that the chromatin at the 4-cell stage obtained a slightly more compressed conformation compared to the 2-cell stage. Previous studies have demonstrated that at the 4-cell stage, more heterochromatin silencing is observed^38^, and domains become more compartmentalized^39^ as the embryo begins to transition out of the totipotent state.

### Section 5 Integrated Pseudo Time Analysis of sc-RT and RNA Recapitulated Developmental Progression

Pseudo time trajectory analyses are often performed on scRNA-seq transcriptome datasets to observe how single cells progress through stem cell differentiation, cancer evolution, and the like^40,41^. With the sc-multi omics protocol, we have access to both DNA and mRNA from each cell. We utilized this depth of information and performed an integrated pseudo time analysis that integrates scRT+ scRNA information per cell. We observed that the integrated trajectory conserved the observed real time developmental progression (**Fig. 5d**). We also observed stage-specific clustering in the final trajectory.

## Discussion

Here, we demonstrated that RT was established at the zygote/1-cell stage before the major-ZGA at the 2-cell stage with conserved patterns observed across zygotes and also between sc-RT datasets. We observed that the replication domains became smaller and more defined as the embryos progressed from the zygote to the 4-cell stage consistent with the establishment of more defined compartments at the 4-cell stage^38,39^. We observed that RT correlated with gene expression in the embryos. At these earliest embryonic stages, late replicating bins had significantly higher gene expression compared to the early bins. The opposite trend has been observed in somatic cells. Unlike somatic cells, mouse embryos until the 4-cell stage are totipotent, have a very short G1 phase (G1 phase is important for establishing stable nuclear organization^42^) and undergo rapid chromatin reorganization. Further, heterochromatin until the 2-cell stage is more ‘liquid-like’ and gains a more silenced conformation beginning at the 4-cell stage^38^. As the embryo transitions from totipotent to pluripotent and then differentiated cell types, the cells gain more defined nuclear organization and topological associated domains (TADs)^39^. These factors can contribute to the difference in RT-RNA trends that we observed in the early embryos.

Further, in previous studies involving somatic cells and global RT-RNA correlation, the transcriptome consisted of constitutively expressed genes which are always early replicating, irrespective of the cell type^8^. These constitutively expressed genes dominated the RT-RNA trend, which is a drawback of the global correlation studies^2,8,43^. When considering only cell-type specific genes for RT-RNA, it was observed that some genes were late replicating and were expressed by switching from early-to-late replication time^43^, not unlike what we observed in our embryos. In this study, by correlating reference ATAC-seq data with the RT and RNA data, we were able to gain insight into multiple levels of organization within the cell. This extensive multilevel dataset can be used to perform network analyses, at the level of individual genes. More importantly, we demonstrated that RT was established before active transcription began in the early embryo. However, it is possible that the maternal mRNAs, which are translated in the zygote^44^, contribute to chromatin organization and defining RT at the zygote stage. Future studies are required to define RT in the absence of all or some of the maternal mRNAs.

## Methods

1. **Isolation of single cells from zygotes, 2-cell, and 4-cell embryos** Mouse zygotes at E0.5 were extracted from super ovulated C57/Bl6 mice. Embryos were pooled from 20 mice per developmental stage analyzed. Zygotes were cultured *in vitro* until 2-cell and 4-cell stages. Zygotes/1-cell embryos were directly transferred into RLT buffer in 96-well plates with 1 zygote/ well. Single cells from 2-cell embryos were extracted 26-28 hours after zygote extraction. Single cells from 4-cell embryos were extracted 38-40 hours after zygote extraction. For 2-cell and 4-cell stages, first the zona pellucida was removed using the Tyrode Acid Solution, then the embryos were incubated in TrypLE: Accutase (1:3) solution for ∼10 minutes until the embryos dissociated into single cells. Single cells were then mouth pipetted into individual wells of a 96-well plate containing RLT buffer. The wells were spun down and stored at -20°C until further processing.
2. **Processing of single cells from embryos using the single cell multiomics protocol** Single cells were isolated from embryos using a mix of Trypsin: Accutase (1:3) for dissociation into single cells, after removing the zona pellucida using Tyrode Acid solution. Single cells from the zygote (n=36), 2-cell (n=42), and 4-cell (n=43) stages were processed using the in-house single cell multiomics protocol. The step-by-step protocol has been uploaded on protocols.io (link in data availability section). Briefly, gDNA and mRNA were extracted, and amplified from each single cell. The separation of gDNA from RNA, and processing of cDNA were adopted from the G&T protocol and Smart-seq2 technique with minor changes^18,19^. The gDNA was processed using an in-house gDNA processing protocol. 22 cycles of cDNA amplification and 22-24 cycles of gDNA amplification was used for the single cells from zygotes, 2-cell, and 4-cell embryos. Quality control of amplified gDNA and cDNA was performed at intermediate steps. Amplified sequences were prepared for sequencing using the Nextera XT DNA Library Prep Kit and each sample was tagged with unique oligo barcodes. All uniquely tagged gDNA and cDNA were pooled together and sequenced. A sequencing depth of ∼2 million reads per gDNA sample and ∼1 million per cDNA sample was sufficient for deducing replication timing and gene expression information respectively. We used the Illumina NextSeq P1 chip with 300 cycles (150 PE).
3. **Analysis of single cell replication timing for zygotes, 2-cell and 4-cell stages** We used the Kronos scRT analysis pipeline^26^ for generating single cell RT binarized heatmaps and pseudo bulk plots for each chromosome genome wide. G1 mESCs (GSE108556) were employed as G1 controls for normalizing gDNA reads of the S-phase embryo cells. We used bin sizes of 200kb for the scRT plots. The RT UMAP was computed using the *umap* R library on the RT values per bin and cell, and the PCA components were computed with the *prcomp* function from the *stats* package.
4. **Analysis of gene expression in zygotes, 2-cell, and 4-cell stages** We used the Seurat v4 pipeline for the single cell RNA analysis and for generating the UMAP, heatmaps, and clusterProfiler for the GO analysis. The pseudo time trajectory analysis was done using Monocle3. Correlation between scRNA seq data generated in this study with reference databases performed for the zygote, 2-cell and 4-cell stages using reference scRNA databases-GSE66582 and GSE45719.
5. **Replication timing and gene expression correlation analyses** Genes with FPKM>1 and expressed in at least 75% of cells per stage were used for the RT-RNA correlation analyses. For generating the post ZGA list for zygote, 2-cell and 4-cell embryos, we utilized genes with (FPKM ≤ 0.5) in oocytes (GSE66582) that were activated (FPKM>1) in the relevant stage of zygote, 2-cell, and 4-cell embryos. Gene-specific RT was calculated from the pseudo bulk genome wide RT plot for each stage. For RT bins values genome-wide, we calculated the probability of gene expression. Slope of correlation between gene expression and RT was plotted for each stage. For the individual cells’ correlation analysis, RT-RNA slopes were calculated within each cell, similar to the strategy used for the pseudo bulk analysis. The values of the slopes were plotted against an individual cell’s progression through S phase.
6. **Genome-wide clustering analysis and replication timing, and gene expression switches per cluster** Genome was divided into 20kb bins and subjected to hierarchical clustering with number of clusters set to 8. The mean RT and gene expression of post-ZGA genes within each cluster was calculated. For comparison with ATAC-seq reference dataset (GSE66581), the mean ATAC score of genes in each cluster was plotted.
7. **Integrated pseudo time trajectory analysis** For integrated analysis, we created a filtered Seurat object. RNA assay with RNA principal components (PCs) was established after normalization and scaling. Similarly, an RT assay with RT PCs was created. RNA and RT UMAPs were produced using multimodal neighbors from the RNA and RT PCAs. Using the top 20 PCs per cell, we generated a UMAP and performed the trajectory inference on the UMAP.

## Data Availability Section

- All code used for the analysis has been uploaded at GitHub link-https://github.com/AnalaShetty1/Embryo-sc-multiomics.git.
- All raw and processed sequencing data generated in this study have been submitted to the NCBI Gene Expression Omnibus (GEO; https://www.ncbi.nlm.nih.gov/geo/)”) under accession number GSE284010.
- The detailed steps for the single cell multiomics protocol are at protocols.io link-DOI: dx.doi.org/10.17504/protocols.io.36wgqdnwyvk5/v1

## Acknowledgements

We thank Dr. Juan Carlos Rivera-Mulia for early discussion and support on this project.

This work is supported by NIH grants R01 DK117286 (CJS), R01 DK117286-03S1 (CJS and WCL), R01 AI173804-01 (CJS and WCL).

Images were created using BioRender https://BioRender.com

## Notes

### Competing Interest Statement

The authors have declared no competing interest.

