## Extended Figures for "Novel Single-Cell Multiomics Approach to Analyze Replication Timing and Gene Expression in Mouse Preimplantation Embryos"

### Extended Data Figures-

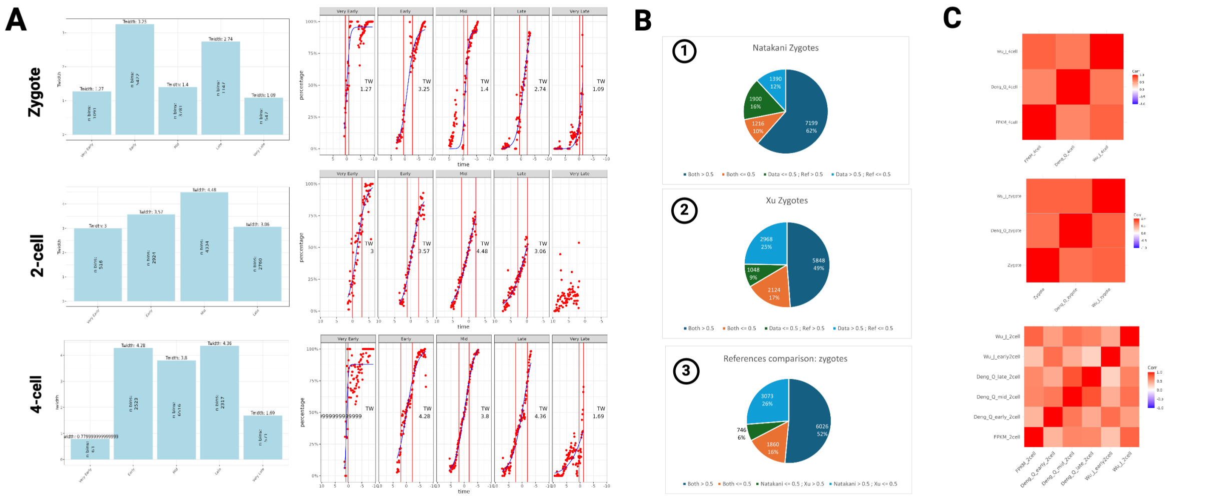


*Extended Data Figure 1:* **(a)** Twidth plots (per stage trends on left, and per cell trends on right) that depict the number of bins replicated in different phases of the cell cycle for zygote, 2-cell, and 4-cell embryonic stages. **(b)** Bins that match and mismatch in Zygote RT profiles between (1) This study's zygote dataset and Nakatani reference dataset (GSE218365); (2) this study's zygote dataset and Xu reference dataset (PRJNA874697); and (3) between Nakatani and Xu reference datasets. **(c)** Correlation between this study's zygote, 2-cell, and 4-cell gene expression datasets with reference datasets (GSE66582 and GSE45719).

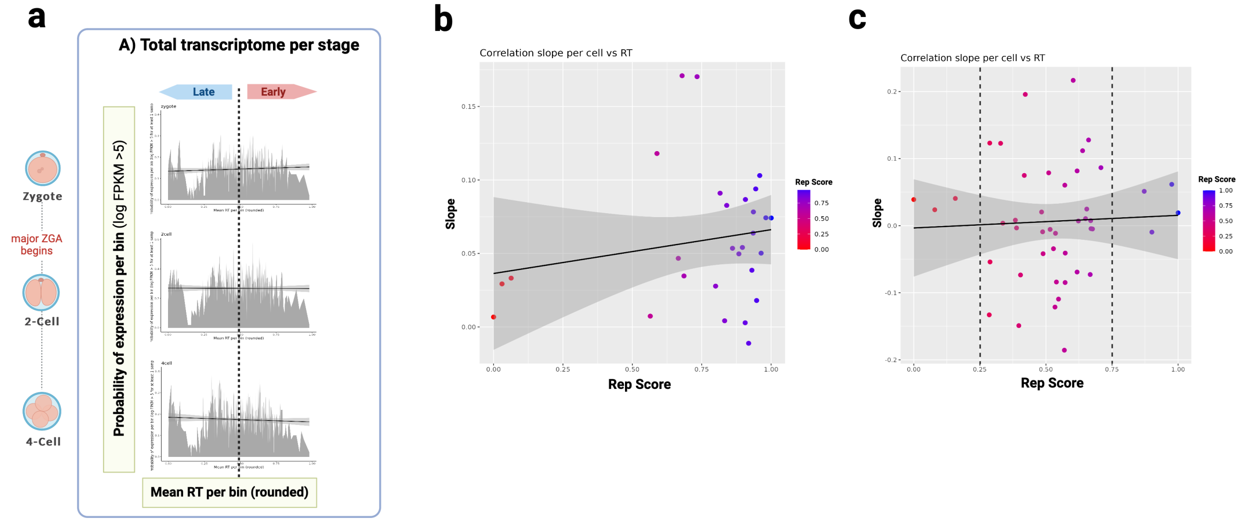


*Extended Data Figure 2:* **(a)** RT and gene expression pseudo bulk analysis for zygote, 2-cell, and 4-cell embryonic stages using total transcriptome per stage. This includes the maternal inherited transcripts. Replication timing and gene expression correlation for individual cells from **(b)** Zygote and **(c)** 4-cell stage. For each cell, the RT-gene expression correlation slopes were calculated and plotted on the Y axis. The Rep Score for each cell, that is, the percent of genome replicated, was plotted on the X axis.

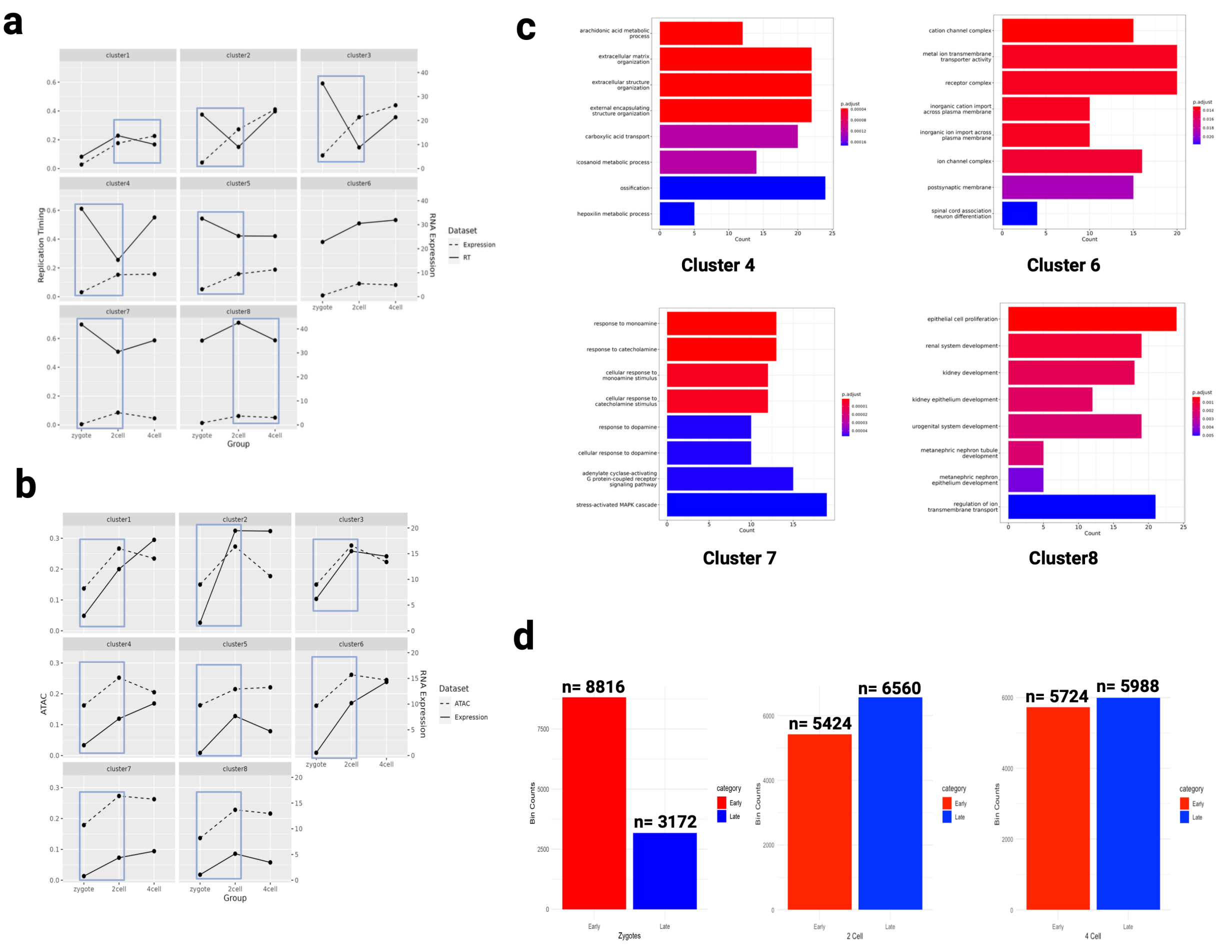


*Extended Data Fig. 3:* **(a)** Trends of changes in RT and gene expression for all clusters 1-8 from Figure 5a using total transcriptome across all clusters. Blue boxes highlight trends of late-to-early RT changes and corresponding lower-to-higher gene expression changes. **(b)** Trends of changes in gene expression and chromosome accessibility (ATAC-seq values) for all clusters 1-8 from Figure 5A. Blue boxes highlight trends of lower-to-higher gene expression changes and corresponding closed-to-open chromatin changes (higher ATAC score). **(c)** Gene ontology analysis of clusters from Figure 5a. **(d)** Genome-wide early and late bin counts from pseudo bulk RT profiles of Zygote, 2-cell, and 4-cell embryos.
